# GABA plays a key role in plant acclimation to a combination of high light and heat stress

**DOI:** 10.1101/2021.02.13.431103

**Authors:** Damián Balfagón, Aurelio Gómez-Cadenas, José L. Rambla, Antonio Granell, Carlos de Ollas, Ron Mittler, Sara I Zandalinas

## Abstract

Plants are frequently subjected to different combinations of abiotic stresses, such as high light intensity and elevated temperatures. These environmental conditions pose an important threat to agriculture production, affecting photosynthesis and decreasing yield. Metabolic responses of plants, such as alterations in carbohydrates and amino acid fluxes, play a key role in the successful acclimation of plants to different abiotic stresses, directing resources towards stress responses and suppressing growth. Here we show that the primary metabolic response of *Arabidopsis thaliana* plants to high light or heat stress is different than that of plants subjected to a combination of high light and heat stress. We further demonstrate that a combination of high light and heat stress results in a unique metabolic response that includes increased accumulation of sugars and amino acids, coupled with decreased levels of metabolites participating in the tricarboxylic acid (TCA) cycle. Among the amino acids exclusively accumulated during a combination of high light and heat stress, we identified the non-proteinogenic amino acid γ-aminobutyric acid (GABA). Analysis of different mutants deficient in GABA biosynthesis, in particular two independent alleles of glutamate decarboxylase 3 (*gad3*), reveal that GABA plays a key role in the acclimation of plants to a combination of high light and heat stress. Taken together, our findings identify a new role for GABA in regulating plant responses to stress combination.

**One sentence summary:** The non-proteinogenic amino acid γ-aminobutyric acid (GABA) is required for plant acclimation to a combination of high light and heat stress in Arabidopsis.

## INTRODUCTION

Plants growing under natural conditions are exposed to different abiotic and biotic stresses that impact plant growth and development. Among these, high light intensities that exceed the plant photosynthetic capacity often occur in native habitats (Ort, 2001; Roeber et al., 2020). Because light plays a key role in the life of photosynthetic organisms, plants evolved many different acclimation and adaptation mechanisms to counteract the effect of high light stress, including paraheliotropic movements, pathways for adjusting the size of the antenna complexes, quenching mechanisms, and pathways to scavenge excess reactive oxygen species (ROS; Asada, 2006; Li et al., 2009; Dietz, 2015). The excess excitation energy produced at the antennas of the photosynthetic apparatus during high light stress is potentially dangerous and could lead to irreversible damage to the reaction centers. Consequently, a sustained decrease in efficiency and electron transport rates could occur, leading to photoinhibition (Ruban, 2015). In addition to high light stress, heat stress can also compromise PSII electron transport due to the increase in fluidity of the thylakoid membranes, dislodging of PSII light harvesting complexes and decreasing the integrity of PSII (Mathur et al., 2014). Moreover, because CO_2_ fixation is dependent on stomatal regulation and temperature, high light stress may cause a more severe hazard to plants when combined with other stresses that already limit the rates of CO_2_ fixation (Mittler, 2006; Roeber et al., 2020). It was recently reported that a combination of high light and heat stress displayed unique transcriptomic, physiological and hormonal responses in *Arabidopsis thaliana* plants (Balfagón et al., 2019). In addition, this abiotic stress combination was found to have a severe impact on PSII performance and to decrease the ability of plants to repair PSII (Balfagón et al., 2019). Lipophilic antioxidant molecules were previously shown to contribute to the protection of PSII against photodamage and enhance tolerance of tomato plants to high light and heat stress combination (Spicher et al., 2017). In sunflower, changes in the steady-state level of transcripts associated with energy metabolism were found in response to this stress combination (Hewezi et al., 2008). The specific physiological and molecular responses observed in different plant species in response to a combination of high light and heat stress (Hewezi et al., 2008; Spicher et al., 2017; Balfagón et al., 2019), could in turn lead to changes in plant metabolism that would minimize stress-induced damages (Balfagón et al., 2020).

Metabolites play an essential role in plant growth and development, as well as modulate different environmental responses of plants. The plant metabolome consists of a wide variety of low molecular weight compounds with many different biological functions, such as carbohydrates that are direct products of photosynthesis and substrates of energy metabolism; tricarboxylic acid (TCA) cycle intermediates; and amino acids involved in protein synthesis and/or other cellular processes such as osmotic readjustments. Increased levels of different polar compounds in plants subjected to different abiotic stresses, including drought, salinity, high light, and extreme temperatures, are thought to play a key role in plant acclimation (Kaplan et al., 2004; Cramer et al., 2007; Maruyama et al., 2009; Caldana et al., 2011). For example, under osmotic stress, TCA cycle, gluconeogenesis and photorespiration are activated to increase glucose, malate and proline levels in order to cope with ROS production and photoinhibition (Cramer et al., 2007). A comparative metabolite analysis of Arabidopsis plants responding to heat or cold shock suggested that a metabolic network consisting of proline, monosaccharides (glucose and fructose), galactinol and raffinose has an important role in tolerance to temperature stress (Kaplan et al., 2004; Urano et al., 2010). Rizhsky et al. (2004) reported that different sugars and amino acids could play a key role in the response of Arabidopsis plants to a combination of drought and heat stress. A study in citrus plants subjected to a combination of drought and heat stress further revealed that the ability of a tolerant citrus genotype to retain a high photosynthetic activity and to cope with oxidative stress was directly linked to its ability to maintain primary metabolic activity (Zandalinas et al., 2016). Moreover, metabolomic analysis of maize plants subjected to drought, heat, and their combination revealed a direct relationship between metabolism and grain yield, highlighting the importance of photorespiration and raffinose family oligosaccharide metabolism for grain yield under drought conditions (Obata et al., 2015).

In general, abiotic stress conditions result in the accumulation of free amino acids in different plants (*e.g.,* Rizhsky et al., 2004; Lugan et al., 2010; Aleksza et al., 2017; Huang and Jander, 2017; Batista-Silva et al., 2019). Several amino acids can act as precursors for the synthesis of secondary metabolites and signaling molecules. For example, polyamines are derived from arginine (Alcázar et al., 2010), and the plant hormone ethylene is synthesized from methionine (Amir, 2010). In addition, a wide range of secondary metabolites with different biological functions are derived from the aromatic amino acids phenylalanine, tyrosine and tryptophan, or from intermediates of their biosynthesis pathways (Tzin and Galili, 2010).

To dissect different primary metabolic responses and to identify promising metabolic markers for a combination of high light and heat stress in plants, we studied the effect of this stress combination on the levels of different primary metabolites in *Arabidopsis thaliana* plants. Both high light and heat stress conditions impacted PSII performance when occurring individually, and their combination displayed unique transcriptomic and physiological responses in plants (Balfagón et al., 2019). We therefore hypothesized that this stress combination would have a unique metabolomic response, leading to the accumulation of metabolites unique to the state of stress combination. Our findings indicate that the primary metabolic response of Arabidopsis plants to a combination of high light and heat stress is different than that of plants subjected to high light or heat stress. We further identified γ-aminobutyric acid (GABA) as a metabolite that specifically accumulates in plants during a combination of high light and heat stress. Using different mutants deficient in GABA accumulation, we further reveal that GABA plays a key role in the acclimation of plants to this stress combination.

## RESULTS

### Physiological responses of Arabidopsis plants to high light, heat stress and their combination

To study the physiological responses of Arabidopsis plants to high light, heat stress and their combination, we subjected wild-type (Col-0) plants to high light intensity (600 μmol m^−2^ s^−1^; HL), high temperature (42°C; HS), or to the combination of HL and HS (600 μmol m^−2^ s^−1^ and 42°C; HL+HS) for 7 hours. Control (CT) plants were maintained at 50 μmol m^−2^ s^−1^ and 23°C during the entire experimental period (Supplemental Fig. S1; Balfagón et al., 2019). Gas exchange parameters, including leaf photosynthetic rate (A), transpiration (E) and stomatal conductance (gs), were determined in Arabidopsis plants subjected to HL, HS and HL+HS (Fig. 1). Photosynthesis, transpiration, and stomatal conductance significantly decreased following the application of HL compared to CT values. These results are in agreement with previous reports showing stomata to close during light stress (Devireddy et al., 2018; Balfagón et al., 2019), limiting transpiration and negatively affecting photosynthetic rates. In contrast, the application of HS increased transpiration and stomatal conductance, maintaining stomata open to cool down the leaf surface via transpiration (in agreement with stomatal aperture measurements; Balfagón et al., 2019). However, HS did not affect photosynthesis compared to CT. Interestingly, the HL+HS combination induced a significant decrease in photosynthesis compared to CT, whereas transpiration and stomatal conductance dramatically increased by about 8-fold compared to CT, or 4-fold compared to HS (Fig. 1). These findings demonstrate that although leaf temperature and stomatal aperture were similar between plants subject to HS or HL+HS (Balfagón et al., 2019), compared to plants subjected to HS, transpiration and stomatal conductance were much higher in plants subjected to the stress combination (HL+HS; Fig. 1). Stomatal aperture measurements and transpiration rates may therefore not always correlate with each other, and it may take higher transpiration rates to cool a leaf during HL+HS combination, potentially a result of heat generated due to dissipation of excess light energy by non-photochemical quenching (NPQ) and/or other protective processes (Czarnocka and Karpiński, 2018; Murchie and Ruban, 2020). Taken together, the results presented in Fig. 1 suggest that during HL+HS combination, HS-associated transpiration and stomatal conductance responses prevailed over those induced by HL (stomatal closure), and that reduction in leaf temperature was more important for plants subjected to HL+HS, than HL-induced stomatal closure that could minimize water loss (Fig. 1; Balfagón et al., 2019).

**Fig. 1.**
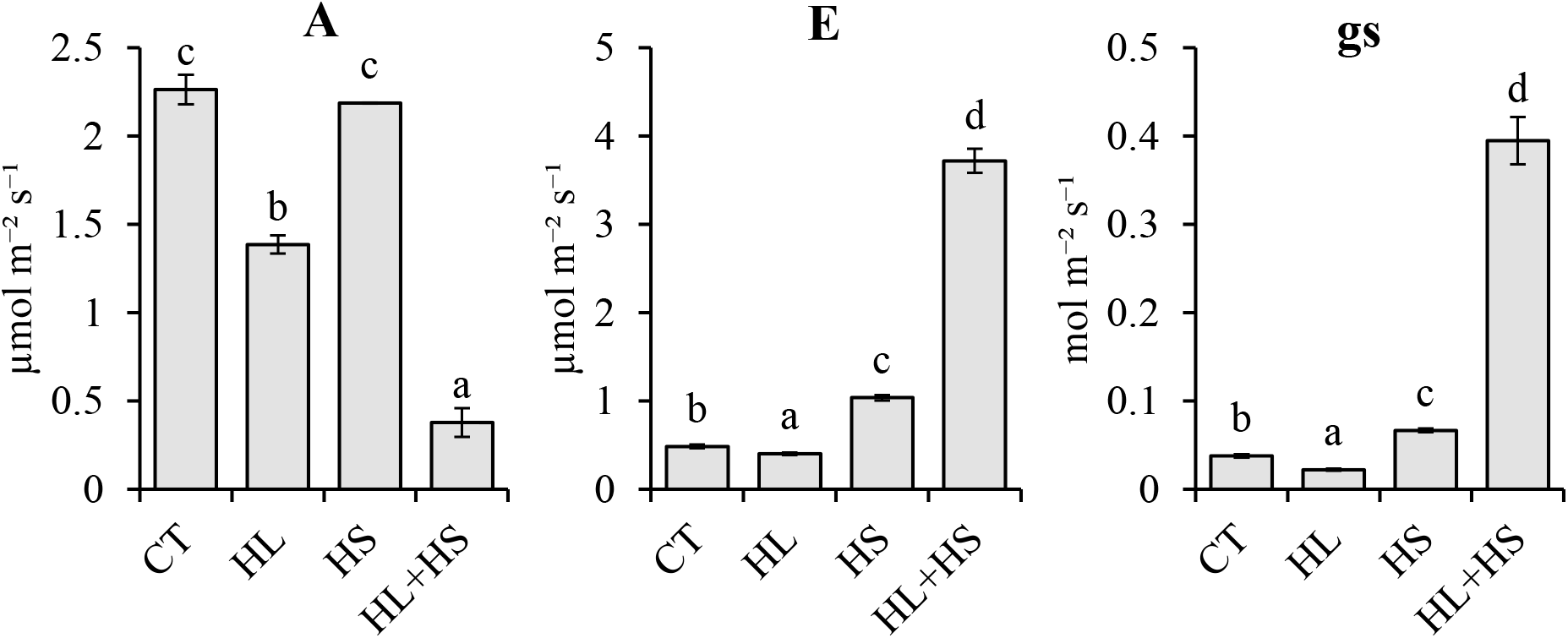
Physiological measurements of Arabidopsis plants subjected to high light, heat stress and their combination. Leaf photosynthetic rate (A), transpiration (E), and stomatal conductance (gs) of Col-0 plants subjected to high light (HL), heat stress (HS) and the combination of HL and HS (HL+HS). Error bars represent SD (N=9). Different letters denote statistical significance at p < 0.05. *Abbreviations used*: A, photosynthetic rate; E, transpiration; gs, stomatal conductance; CT, control; HL, high light; HS, heat stress; HL+HS, a combination of high light and heat stress.

### Metabolomic responses of Arabidopsis plants to high light, heat stress and their combination

To study the accumulation of stress-associated metabolites in Arabidopsis plants subjected to HL, HS or HL+HS, a gas chromatography-mass spectrometric (GC-MS) analysis of polar compounds extracted from leaves of plants subjected to the different stresses was performed (Supplemental Fig. S1). Principal Component Analysis (PCA) revealed that the main source of variation in the data was due to metabolic changes associated with the stress combination, as the first principal component, accounting for 56.6% of total variance, was defined by the characteristic profile of HL+HS samples. In turn, principal component 3, explaining 11.5% of total variation, clearly separated the samples based on the metabolic profile of plants subjected to HS (Fig. 2A). Analysis of variance revealed a total of 25 polar metabolites with levels significantly altered in response to HL (21 and 4 over- and under-accumulated, respectively), levels of 23 metabolites significantly altered in response to HS (19 and 4 over- and under-accumulated, respectively), and levels of 38 metabolites changed compared to CT under HL+HS (28 and 10 over- and under-accumulated, respectively) (Fig. 2B; Table 1). Moreover, of the 28 metabolites with levels significantly elevated in response to HL+HS, 3 metabolites (10.7%) were common with HL-induced metabolites, other 3 metabolites (10.7%) were common with HS-induced metabolites, and 7 metabolites (25.0%) were found to be specifically accumulated in response to HL+HS. Similarly, levels of 1 metabolite (10.0%) commonly decreased in response to either HL or HS, and levels of 7 metabolites (70.0%) were reduced in response to HL+HS (Fig. 2B). These results indicated that a substantial portion of polar metabolites with altered levels in plants subjected to HL+HS was specific for the stress combination. As shown in Table 1, metabolites that exclusively accumulated in plants in response to HL+HS were glycerol, succinic acid, GABA, rhamnose, arginine, gluconic acid and tyrosine. In contrast, levels of threonic acid, urea, fumaric acid, nicotinic acid, citric acid, pyroglutamic acid and putrescine specifically decreased in response to HL+HS (Table 1).

**Fig. 2.**
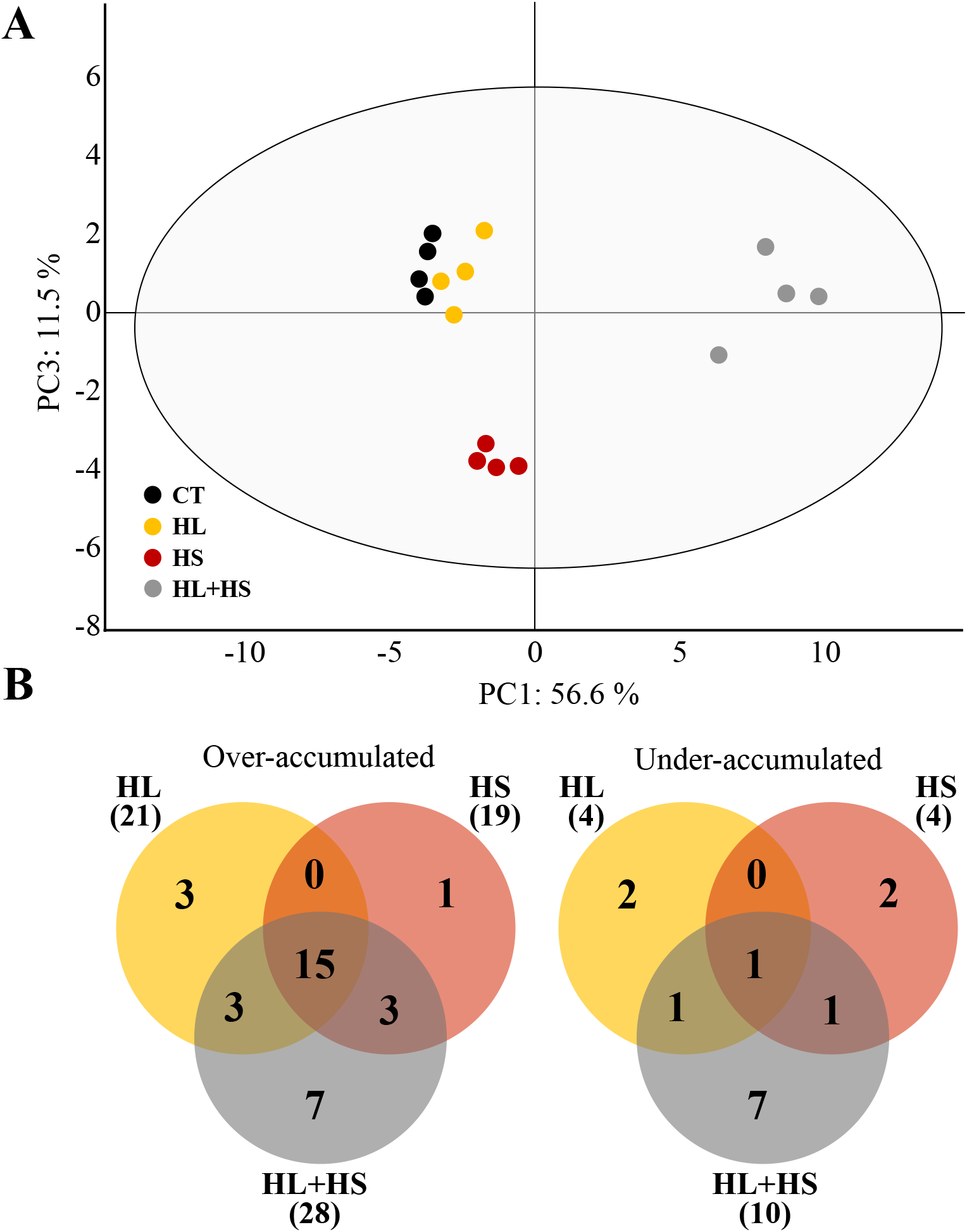
Metabolic analysis of Arabidopsis Col-0 plants subjected to high light, heat stress and their combination. (A) Principal Component Analysis (PCA) score plot of metabolite profiles obtained from control Col-0 plants (CT), and Col-0 plants subjected to high light (HL), heat stress (HS) or a combination of HL and HS (HL+HS). (B) Venn diagrams showing the overlap between metabolites over-accumulated (left) or under-accumulated (right) in response to HL, HS and HL+HS combination. *Abbreviations used*: CT, control; HL, high light; HS, heat stress; HL+HS, a combination of high light and heat stress.

**Table 1.** List of metabolites over- and under-accumulated in Col-0 plants subjected to high light (HL), heat stress (HS) and a combination of HL and HS (HL+HS). Values represent fold changes compared to control. Bold values represent fold changes > 10 for over-accumulated metabolites, and fold changes < 0.5 for under-accumulated metabolites. Metabolites shown are all significant (N=4, t-test, p < 0.05; see Tables S1, S3). *Abbreviations used*: GABA, γ-aminobutyric acid; HL, high light; HS, heat stress; HL+HS, a combination of high light and heat stress.

| Stress | Metabolite | Over-accumulated<br>(Fold change) |  |  | Metabolite | Under-accumulated<br>(Fold change) |  |  |
| --- | --- | --- | --- | --- | --- | --- | --- | --- |
|  |  | HL | HS | HL+HS |  | HL | HS | HL+HS |
| HL | Fumaric acid | 1.90 |  |  | GABA | 0.65 |  |  |
|  | Malic acid | 2.60 |  |  | Rhamnose | 0.69 |  |  |
| | α-ketoglutaric acid | 2.11 | | | | | | |
| HL &<br>HL+HS | Proline | **10.35** |  | **27.35** | Glutamic acid | 0.74 |  | **0.36** |
|  | Methionine | 1.81 |  | 2.37 |  |  |  |  |
|  | Threolose | 2.13 |  | 3.92 |  |  |  |  |
| HS | Putrescine | 1.49 |  |  | Methionine |  | 0.64 |  |
|  |  |  |  |  | Arginine |  | 0.52 |  |
| HS &<br>HL+HS | Asparagine |  | 3.18 | 2.71 | Myoinositol |  | 0.53 | **0.44** |
|  | Lysine |  | 2.57 | **23.90** |  |  |  |  |
|  | Tryptophan |  | 1.95 | **16.72** |  |  |  |  |
| HL+HS | Glycerol |  |  | 2.48 | Threonic acid |  |  | 0.69 |
|  | Succinic acid |  |  | 2.33 | Urea |  |  | 0.55 |
|  | GABA |  |  | **59.43** | Fumaric acid |  |  | 0.58 |
|  | Rhamnose |  |  | 2.32 | Nicotinic acid |  |  | 0.60 |
|  | Arginine |  |  | 1.31 | Citric acid |  |  | **0.47** |
|  | Gluconic acid |  |  | 2.32 | Pyroglutamic acid |  |  | 0.68 |
|  | Tyrosine |  |  | 2.11 | Putrescine |  |  | **0.38** |
| HL & HS &<br>HL+HS | Alanine | **43.61** | **287.70** | **593.02** | Aspartic acid | 0.62 | 0.61 | **0.25** |
|  | Valine | **12.35** | **16.14** | **73.63** |  |  |  |  |
|  | Leucine | 7.82 | 8.10 | **174.49** |  |  |  |  |
|  | Isoleucine | 7.41 | **12.56** | **148.53** |  |  |  |  |
|  | Glycine | 26.28 | 9.14 | **12.78** |  |  |  |  |
|  | Threonine | 2.86 | 2.75 | 3.21 |  |  |  |  |
|  | Erythritol | 1.48 | 1.74 | 3.86 |  |  |  |  |
|  | 4-hydroxyproline | 1.54 | 1.95 | 2.05 |  |  |  |  |
|  | Phenylalanine | 1.78 | 3.64 | **15.04** |  |  |  |  |
|  | Glutamine | 6.64 | 3.56 | **15.18** |  |  |  |  |
|  | Fructose | 1.68 | 1.93 | 7.73 |  |  |  |  |
|  | Glucose | 2.77 | 3.31 | 9.42 |  |  |  |  |
|  | Sucrose | 1.50 | 2.06 | 1.69 |  |  |  |  |
|  | Maltose | 3.72 | 1.83 | **50.28** |  |  |  |  |
|  | Raffinose | 3.07 | **42.54** | **31.74** |  |  |  |  |

### The impact of high light and heat stress combination on sugar metabolism, TCA cycle intermediates, and amino acid levels

Further analysis of metabolites involved in glycolysis, TCA cycle and amino acid biosynthesis during stress combination, was conducted (Fig. 3). The soluble sugars glucose and fructose, as well as raffinose and maltose strongly accumulated in response to HL+HS whereas their accumulation, in general, was less pronounced in response to HL or HS. In addition, trehalose and erythritol accumulated in response to the different treatments and particularly during HL+HS. In contrast, sucrose, the major form of carbohydrates transported from photosynthetically active tissues, slightly increased in its level in response to the individual and combined stresses (Fig. 3A; Table 1; Supplemental Table S1). Analysis of TCA-cycle intermediates revealed that HL+HS perturbed the TCA cycle and reduced the levels of the TCA-cycle-derived amino acids aspartate and glutamate. Aromatic amino acids are synthesized in plants through the shikimate pathway. In our study, levels of tryptophan and phenylalanine significantly increased during HL+HS, whereas individual stresses had a marginal effect on their accumulation. In addition, tyrosine was also accumulated under HL+HS combination, while no change in its levels was found under HL or HS. Amino acids synthesized from pyruvate including alanine, leucine, valine, and isoleucine significantly accumulated under all stress conditions, although more noticeably under HL+HS conditions (Fig. 3A; Table 1; Supplemental Table S1). The reduction in aspartate levels under HL+HS was accompanied by accumulation of asparagine, methionine, threonine, and especially lysine, whose accumulation was especially high in response to HL+HS (Fig. 3A; Table 1; Supplemental Table S1). Analysis of the expression of genes encoding for enzymes that participate in different reactions of the TCA cycle revealed different patterns of transcript accumulation among the individual and combined stresses (Fig. 3B; Supplemental Table S2). In general, although TCA-related metabolites were suppressed in response to a combination of HL+HS, expression of transcripts encoding TCA cycle-related enzymes increased in response to the stress combination (Fig. 3B; Supplemental Table S2), possibly as a response to counteract the low metabolite accumulation.

**Fig. 3.**
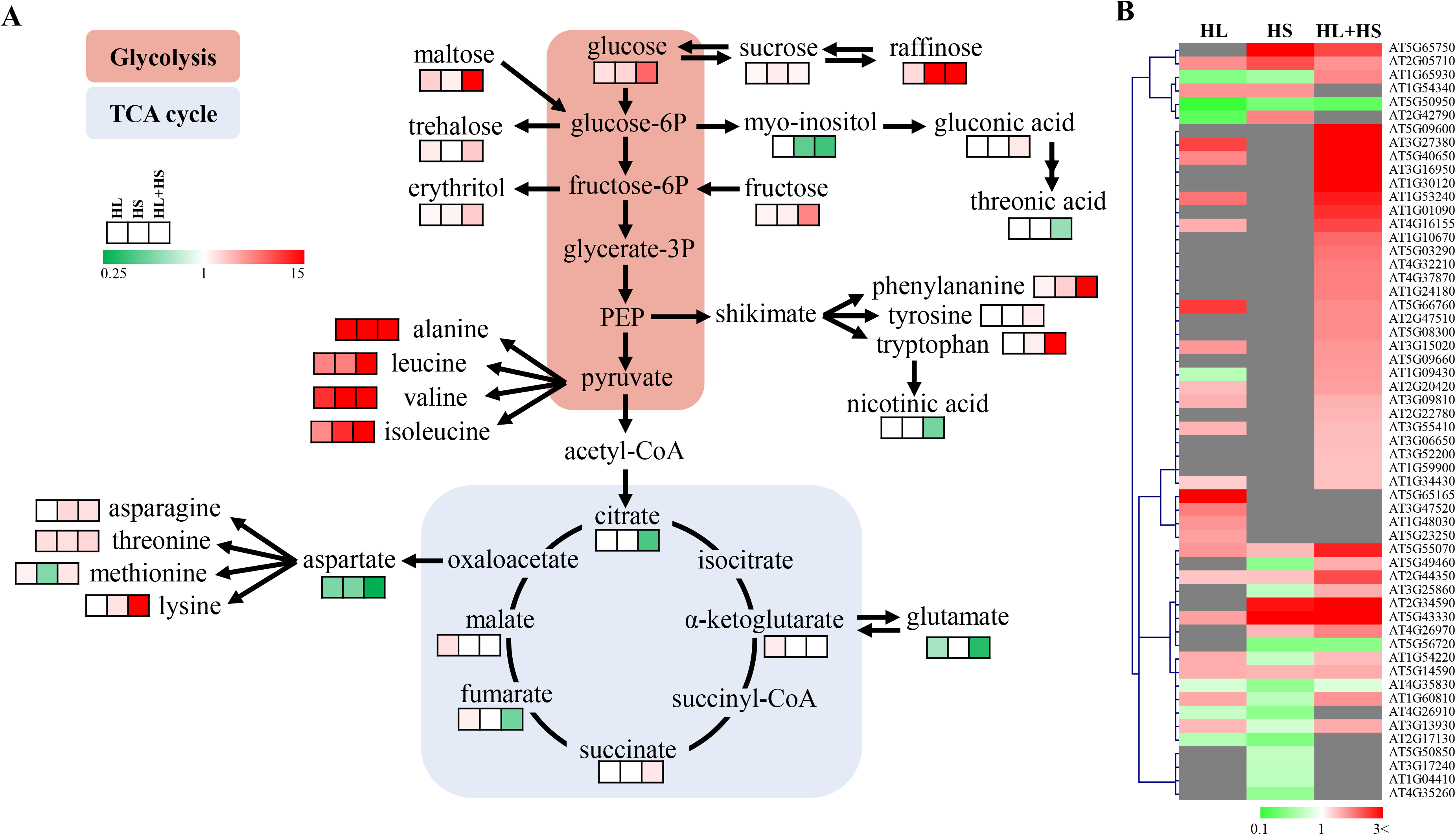
Levels of amino acids and metabolites involved in glycolysis and TCA cycle in Arabidopsis plants subjected to high light, heat stress and their combination. (A) Levels of metabolites participating in glycolysis, TCA cycle, and amino acid metabolism in Col-0 plants subjected to high light (HL), heat stress (HS) or a combination of HL and HS (HL+HS). Significant metabolite levels (p < 0.05) are expressed as fold change compared to control conditions and are shown as a color scale (Table S1). Non-significant accumulation compared to control is shown in white. (B) Heat map showing the expression levels of transcripts involved in TCA cycle in Col-0 plants subjected HL, HS and HL+HS combination. Significant transcript levels (p < 0.05) are expressed as fold change compared to control conditions and are shown as a color scale. Non-significant expression levels compared to control are shown in grey. Transcript expression data was obtained from the RNA-Seq analysis conducted by Balfagón et al. (2019) (Table S2). *Abbreviations used*: HL, high light; HS, heat stress; HL+HS, a combination of high light and heat stress; PEP, phosphoenolpyruvate.

### Impact of a combination of high light and heat stress on glutamate metabolism

Glutamate has a central role in amino acid metabolism in plants, and is also a substrate for the synthesis of arginine, ornithine, proline, glutamine and GABA (Forde and Lea, 2007). As shown in Fig. 4A and Supplemental Table S3, the observed decline in glutamate levels in response to HL+HS was associated with proline, glutamine, and GABA accumulation. In contrast, levels of arginine and urea, as well as levels of the polyamine putrescine, decreased or did not change in response to the application of stress (Fig. 4A; Supplemental Table S3). It was reported that under abiotic stress conditions, oxidation of putrescine contributes to GABA production (Shelp et al., 2012), suggesting that the specific decrease in putrescine under HL+HS conditions could lead to GABA accumulation in response to this stress combination. Indeed, as shown in Fig. 4A and Supplemental Table S3, GABA accumulated exclusively in response to HL+HS. To further dissect GABA metabolism in plants in response to a combination of HL and HS, we analyzed the expression of transcripts involved in GABA biosynthesis (*GAD1*, *GAD2*, *GAD3* and *GAD4*) as well as the expression of transcripts related to GABA catabolism (*POP2* and *ALDH5F1*; using RNA-Seq data obtain by Balfagón et al., 2019). As shown in Fig. 4B and Supplemental Table S4, the expression of *GAD1* and especially *GAD3* remarkably increased only in response to HL+HS. In contrast, expression of *GAD2* was repressed in response to HS and transcript accumulation of *GAD4* slightly increased in plants subjected to the individual HL or HS treatments. The expression of *POP2* decreased in response to HL and HL+HS and all stresses reduced the expression of *ALDH5F1* (Fig. 4B; Supplemental Table S4). The findings presented in Fig. 4 and Supplemental Table S4 suggest therefore a possible role for GABA in regulating plant responses to HL+HS.

**Fig. 4.**
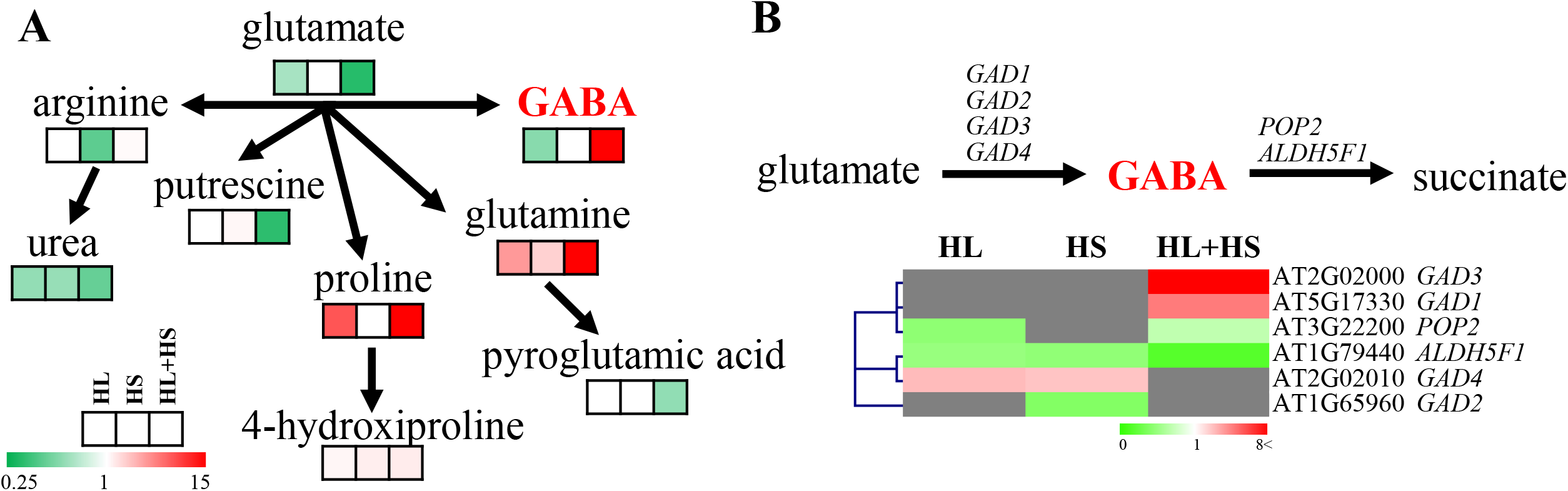
Glutamate metabolism in Arabidopsis plants subjected to high light, heat stress and their combination. (A) Level of metabolites involved in glutamate metabolism in Col-0 plants subjected to high light (HL), heat stress (HS) or a combination of HL and HS (HL+HS). Significant metabolite levels (p < 0.05) are expressed as fold change compared to control conditions and are shown as a color scale (Table S3). Non-significant accumulation compared to controls are shown in white. (B) Heat map showing the expression levels of transcripts involved in GABA metabolism in Col-0 plants subjected HL, HS and HL+HS combination. Non-significant expression levels compared to controls are shown in grey. Transcript expression data was obtained from the RNA-Seq analysis conducted by Balfagón et al. (2019) (Table S4). *Abbreviations used*: ALDH5F1, aldehyde dehydrogenase 5F1; GABA, γ-aminobutyric acid; GAD, glutamate decarboxylase; HL, high light; HS, heat stress; HL+HS, a combination of high light and heat stress; POP2, pollen-pistil incompatibility 2.

### Involvement of GABA in plant tolerance to the combination of high light and heat stress

To further study the role of GABA in the response of plants to HL+HS, we analyzed the response of two independent lines of the GABA-deficient mutant *gad3* (SALK_138534C and SALK_033307C) to HL, HS and HL+HS combination (Fig. 5A). Accumulation of GABA was repressed or did not change in *gad3* mutants subjected to HL or HS, as well as in wild type plants in response to HL. In contrast, HL+HS induced a pronounced increase in GABA levels in Col-0 plants, whereas both *gad3* mutants slightly accumulated GABA, probably due to the action of *GAD1* (Figs. 4B, 5B). The reduced accumulation of GABA in *gad3* plants in response to HL+HS compared to Col-0 (Fig. 5B) was accompanied by a significant decrease in the survival of *gad3* mutants in response to a combination of HL+HS (Fig. 5A, C). Whereas all *gad3* plants survived the individual HL or HS, the survival rate of both *gad3* mutants subjected to HL+HS combination decreased by about 40%. Furthermore, analysis of Leaf Damage Index (LDI; Balfagón et al., 2019) of Col-0 and *gad3* mutants subjected to the different stresses (Fig. 5D) revealed that HL+HS negatively impacted leaf appearance of both *gad3* lines, with 51.2% and 50.2% of leaves dead, 28.2% and 32.7% of leaves injured, and only 20.6% and 16.9% of leaves appearing healthy, in SALK_138534C and SALK_033307C, respectively (Fig. 5D). Compared to Col-0 plants, *gad3* mutants were therefore more sensitive to HL+HS combination.

**Fig. 5.**
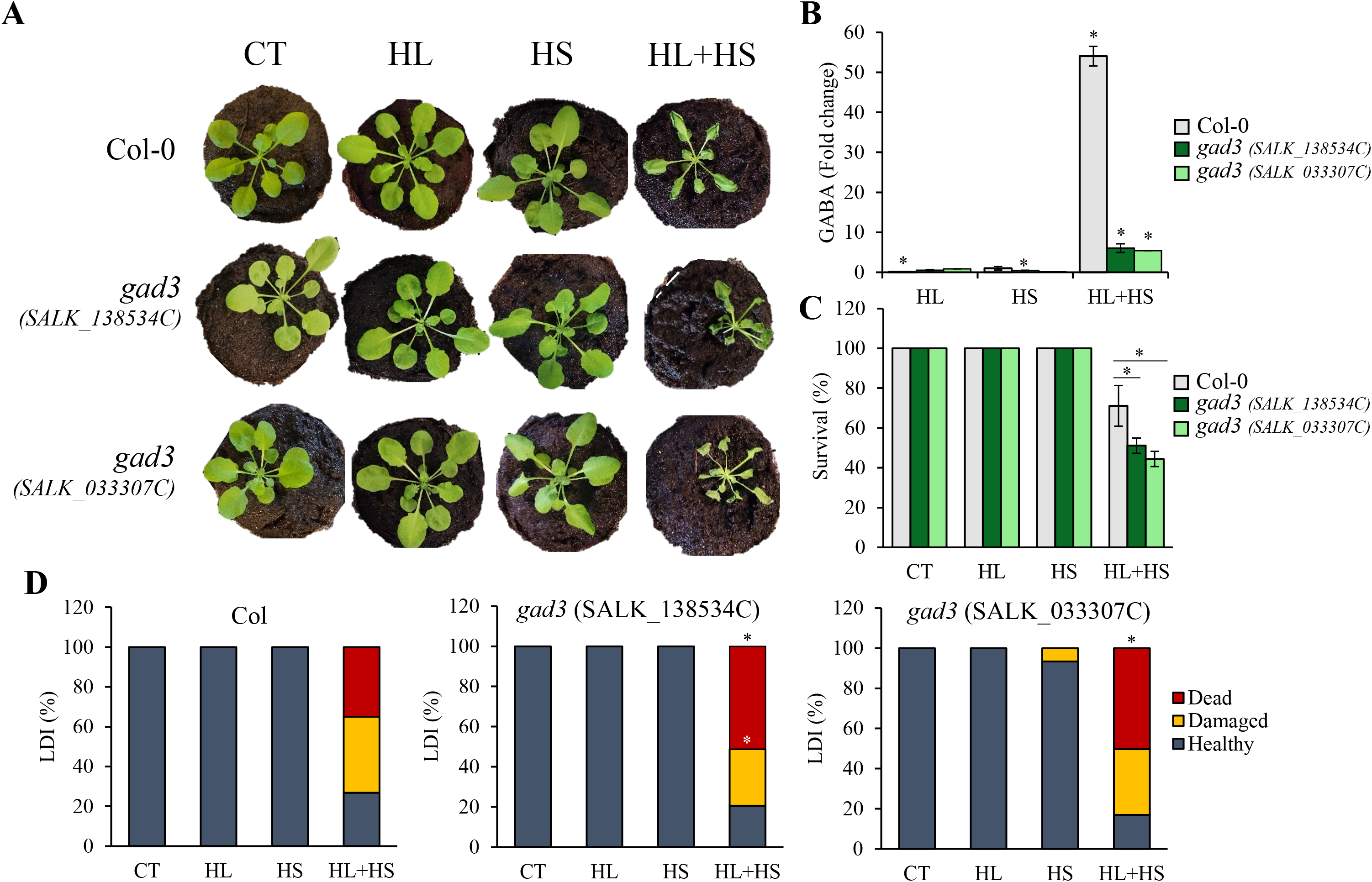
Involvement of GABA in the response of Arabidopsis plants to high light, heat stress and their combination. (A) Representative images of Col-0 and the GABA mutant *gad3* (two independent knockout lines; SALK_138534C and SALK_033307C) subjected to high light (HL), heat stress (HS) and a combination of HL and HS (HL+HS). (B) Levels of GABA in Col-0 and the GABA knockout mutant *gad3* (two independent lines) subjected to HL, HS and HL+HS combination. (C) Survival of Col-0 and the GABA mutant *gad3* (two independent lines) subjected to HL, HS and HL+HS combination. (D) Leaf Damage Index (LDI) of Col-0 and the GABA mutant *gad3* (two independent lines) subjected to HL, HS and HL+HS combination. Asterisks denote Student’s t-test significance at p < 0.05 compared to wild type (C) or to control (B and D). Error bars represent SD. *Abbreviations used*: CT, control; GAD, glutamate decarboxylase; HL, high light; HS, heat stress; HL+HS, a combination of high light and heat stress; LDI, Leaf Damage Index.

## DISCUSSION

The ability of plants to sense and react to different adverse conditions in their environment by modulating physiological responses, gene expression and metabolism, is crucial for plant adaptation and survival during stress. Due to the frequent occurrence of HL+HS combination in nature, and its impact on crops (Yamamoto et al., 2008; Suzuki et al., 2014; Roeber et al., 2020), as well as its impact on plant survival (Balfagón et al., 2019), the study of metabolic changes during this stress combination is of particular interest. A recent study of the physiological and transcriptomic responses of Arabidopsis plants subjected to HL, HS and their combination (HL+HS) revealed that the HL+HS combination was accompanied by irreversible damage to PSII, decreased D1 (PsbA) protein levels, enhanced accumulation of the hormones jasmonic acid (JA) and JA-isoleucine (JA-Ile), elevated expression of over 2,200 different transcripts unique to the stress combination, distinctive structural changes to chloroplasts and a decreased survival rate (Balfagón et al., 2019). In the present study, we show that HL+HS combination has a detrimental effect on photosynthetic rates, and that the effects of HS on stomatal responses and transpiration (opening of stomata and increasing transpiration) prevails over the effects of HL (closing of stomata and decreasing transpiration; Balfagón et al., 2019) (Fig. 1). This result is different than the response of plants to a combination of drought and heat stress, in which the effects of drought prevailed over the effects of heat on stomatal regulation (Rizhsky et al., 2002; Rizhsky et al., 2004). The observed decrease in photosynthetic rates under HL+HS (Fig. 1) prompted us to analyze the primary metabolism of plants subjected to this stress combination to unravel specific patterns of sugar, amino acid and polyamine accumulation (Figs. 2–4; Table 1; Supplemental Tables S1-4). Individual and combined HL and HS displayed different polar accumulation patterns (Fig. 2; Table 1), suggesting that the different stress conditions alter the primary metabolism in different ways, reinforcing the idea that metabolic changes due to stress combination are unique and not a mere additive combination of the effects of each individual stress.

In our study, the levels of several metabolites appeared to be correlated with plant sensitivity to HL+HS combination (Balfagón et al., 2019). Plants subjected to this stress combination accumulated sugars such as glucose, fructose, raffinose, maltose and trehalose, whereas the levels of sucrose slightly increased in response to individual and combined stresses (Fig. 3; Table 1; Supplemental Table S1). The source of sugars in plants subjected to a combination of HL and HS is unknown. Taking into consideration that photosynthesis is suppressed in plants subjected to HL+HS combination (Fig. 1), sugars could be synthesized by way of starch degradation, as proposed to occur during a combination of drought and heat stress (Rizhsky et al., 2004). Indeed, high accumulation of maltose, a major sugar associated with starch degradation (Thalmann and Santelia, 2017), and of its derived sugars were observed in HL+HS-stressed plants (Fig. 3; Table 1; Supplemental Table S1). Additional studies are, however, required to examine this possibility. The increased accumulation of sugars participating in glycolysis under HL+HS (Fig. 3; Supplemental Table S1) suggests that this pathway could provide an alternative source of ATP in plants subjected to HL+HS stress combination, to counteract the negative effects of the stress combination on PSII and photosynthetic rates (Fig. 1; Balfagón et al., 2019), as well as to function as compatible solutes (Krasensky and Jonak, 2012; Shaar-Moshe et al., 2019). Moreover, the levels of glycolysis-derived aromatic amino acids produced through the shikimate pathway, tryptophan, phenylalanine and tyrosine, as well as amino acids synthesized from pyruvate, including alanine, leucine, valine and isoleucine significantly increased during a combination of HL+HS (Fig. 3; Table 1; Supplemental Table S1). Although glycolysis appeared to be activated under HL+HS combination, a concomitant activation of the TCA cycle was not observed (Fig. 3; Table 1; Supplemental Table S1), similar to the findings of Shaar-Moshe et al. (2019), demonstrating that organic acids produced by the TCA cycle were reduced under the combination of salinity, drought and heat. Therefore, depletion of metabolites related to the TCA cycle under stress combination could indicate that respiration might be compromised in plants subjected to HL+HS combination. The reduction in oxalacetate-derived aspartate levels under HL+HS conditions was accompanied by an increase in aspartate-related amino acids, especially lysine (Fig. 3; Table 1; Supplemental Table S1). Increased accumulation of amino acids has been shown in plants subjected to different abiotic stresses (*e.g.*, Kaplan et al., 2004; Rizhsky et al., 2004; Kempa et al., 2008; Sanchez et al., 2008; Usadel et al., 2008; Lugan et al., 2010; Krasensky and Jonak, 2012), and could be a result of amino acid biosynthesis and/or enhanced stress-induced protein degradation. In this sense, the higher impact of HL+HS combination on plant physiology and survival (Fig. 1; Balfagón et al., 2019) could lead to an increase in protein degradation and therefore, higher amino acid content. Further studies elucidating this possibility are needed. The decreased levels of TCA-derived glutamate in response to HL and especially in response to HL+HS combination (Figs. 3, 5; Table 1; Supplemental Table S1) was further accompanied by a concomitant decrease in putrescine (Fig. 4; Table 1; Supplemental Table S3). These results indicate that the role of polyamines under HL+HS as osmoprotectants might be marginal, and that other metabolites including sugars (Fig. 3) and/or proline (Fig. 4) could have a key role as osmoprotective elements under this stress combination. In addition, as a compatible solute, proline is involved in the stabilization of proteins and protein complexes in the chloroplast and cytosol, protection of the photosynthetic apparatus and enzymes involved in detoxification of ROS, as well as redox balance stabilization (Szabados and Savouré, 2010). The high accumulation of proline observed in plants subjected to HL+HS could therefore suggest that the stress combination imposes a stronger pressure on plant metabolism, as indicated by the decrease in survival rates and values of LDI (Fig. 5; Balfagón et al., 2019).

Interestingly, GABA levels were specifically elevated in plants subjected to HL+HS combination (Fig. 4A Table 1; Supplemental Table S3), and GABA-deficient mutants (*gad3*) showed a significant decline in their ability to acclimate to this stress combination (Fig. 5), suggesting that GABA could be required for plant acclimation to a combination of high light and heat stress. GABA is a key non-proteinogenic amino acid that displays important physiological functions involved in plant growth regulation (Seifikalhor et al., 2019). Exogenous GABA application to plants was reported to improve tolerance to different environmental stresses (*e.g.,* Shi et al., 2010; Shang et al., 2011; Li et al., 2016; Salvatierra et al., 2016; Priya et al., 2019; Seifikalhor et al., 2020). Furthermore, GABA levels increased in response to different abiotic stress combinations, namely, salt and drought, as well as salt, drought and heat (Shaar-Moshe et al., 2019). GABA was further proposed to act as a signaling molecule during stress (Bouché and Fromm, 2004; Yu et al., 2014; Fromm, 2020). Another potential function of GABA in plant survival during stress could be linked to its role in regulating autophagy (Supplemental Fig. S2; Signorelli et al., 2019), contributing to the recycling of damaged cellular components during stress. Taken together, the results presented here indicate that GABA plays a key role in the response of plants to HL+HS stress combination, and that genes involved in GABA metabolism could be used as potential breeding markers for HL+HS-tolerant crops.

## MATERIALS AND METHODS

### Plant material and growth conditions

*Arabidopsis thaliana* Col-0 (var. Columbia-0) and *gad3* (SALK_138534C and SALK_033307C) plants were grown in peat pellets (Jiffy-7, http://www.jiffygroup.com/) at 23°C under long day growth conditions (12-hour light from 7 AM to 7 PM; 50 μmol m^−2^ s^−1^/12-hour dark from 7 PM to 7 AM).

### Stress treatments

Individual HL and HS, and a combination of HL and HS were applied in parallel as described in (Balfagón et al., 2019) and shown in Supplemental Fig. S1, using 30-day-old *Arabidopsis thaliana* plants (wild type Col-0 and the SALK_138534C and SALK_033307C *gad3* mutants). HL was applied by exposing plants to 600 μmol m^−2^ s^−1^ (Philips, F54T5/TL84/HO/ALTO) at 23°C for 7 hours. HS was applied by subjecting plants to 42°C, 50 μmol m^−2^ s^−1^, for 7 hours. HL+HS combination was performed by simultaneously subjecting plants to 600 μmol m^−2^ s^−1^ of light stress and 42°C for 7 hours. Control plants were maintained at 50 μmol m^−2^ s^−1^, 23°C during the entire experiment. Following the stress treatments, control plants and plants subjected to HL, HS and HL+HS combination were divided into two groups: a group used for sampling leaves for metabolomics analysis as described below; and a group allowed to recover under controlled conditions until flowering time to score for survival. 24 hours following the stress treatments, Leaf Damage Index (LDI; Gallas and Waters, 2015; Balfagón et al., 2019) was recorded (Supplemental Fig. S1). All experiments were carried out at the same time-of-day during the light cycle (from 9 AM to 4 PM) and were repeated at least three times with 30 plants per biological repeat.

### Photosynthetic parameters

Photosynthetic rate (A), transpiration (E) and stomatal conductance (gs) were measured using a LCpro+ portable infrared gas analyzer (ADC BioScientific Ltd., Hoddesdon, UK) under ambient CO_2_ and moisture conditions. Supplemental light was provided by a PAR lamp at 50 or 600 μmol m^−2^ s^−1^ photon flux density, and air flow was set at 150 μmol mol^−1^. After instrument stabilization, at least 10 measurements were taken on three fully expanded leaves of three plants immediately after the 7 hours of individual and combined stress treatments (Supplemental Fig. S1). All experiments were repeated at least three times.

### Determination of primary metabolites

The relative levels of polar metabolites were determined as described in Zanor et al. (2009). Fifteen mg of freeze-dried plant tissue were extracted in 1.4 mL of methanol and 60 μL of an aqueous solution with 0.2 mg mL^−1^ of ribitol, which was used as internal standard. Extraction was performed at 70°C for 15 min in a water bath. The extract was centrifuged at 14,000 rpm for 10 min, and the supernatant was recovered and fractionated adding chloroform and Milli-Q water. After vigorous vortexing and 15 min and centrifugation at 4,000 rpm, 50 μL of the aqueous phase were recovered and dried overnight in a speed-vac. The dry residue was subjected to a double derivatization procedure with methoxyamine hydrochloride (20 mg mL^−1^ in pyridine, Sigma) and *N*-Methyl-*N*-(trimethylsilyl)trifluoroacetamide (Macherey-Nagel). Fatty acid methyl esters (C_8_-C_24_) were added and used as retention index (RI) markers. Analyses were performed on a 6890N gas chromatograph (Agilent Technologies, USA) coupled to a Pegasus 4D TOF mass spectrometer (LECO, St. Joseph, MI). Chromatography was performed with a BPX35 (30 m, 0.32 mm, 0.25 μm) capillary column (SGE Analytical Science Pty Ltd., Australia) with a 2 mL min^−1^ helium flow. Oven programming conditions were as follows: 2 min of isothermal heating at 85°C, followed by a 15°C min^−1^ temperature ramp up to 360°C. Injection temperature was set at 230°C, and the ion source was adjusted to 250°C. Data were acquired after EI ionization at 70 eV, and recorded in the 70–600 m/z range at 20 scans s^−1^. Chromatograms were analyzed by means of the ChromaTOF software. Metabolites were identified by comparison of both mass spectra and retention time with those of pure standards injected under the same conditions. Peak area of each identified compound was normalized to the internal standard area (ribitol) and sample dry weight. All experiments were repeated four times.

### γ-aminobutyric acid quantification

About 5 mg of freeze-dried plant tissue were transferred to a 1.5-mL microcentrifuge tube. Three glass beads and 50 μL of deuterium labelled internal standard γ-aminobutyric acid (GABA-d2) at concentration of 20 ppm was added. Then, 300 μL of cold MeOH:H_2_O (80:20) was added, following sonication in an ultrasound bath with ice for 10 min and centrifugation at 10,000 rpm for 5 min. 250 μL of supernatant were recovered and 250 μL of acetonitrile were added, following filtration through a PTFE 0.2 μm pore size cellulose filter. Final concentration of the deuterated standard (GABA-d2) was 200 ppb. GABA was quantified in plant extracts using a UPLC system (Waters Acquity SDS, Waters Corp., Milford, MA, USA) interfaced to a TQD triple quadrupole (Micromass Ltd, Manchester, UK) mass spectrometer through an orthogonal Z-spray electrospray ion source. Separations were carried out on 2.1 mm × 150 mm ACQUITY UPLC 1.7 m BEH amide Column using a linear gradient of (A) acetonitrile-water 95:5 (v:v), 0.1% ammonium formate and (B) acetonitrile-water 2:98 (v:v), 0.1% ammonium formate at a flow rate of 300 μL min^−1^. Chromatographic run started at 0% B; after 1 min a linear gradient increased A to 75% for 3 min; finally, mobile phase composition returned to the initial conditions for 2 min. Transitions for GABA (104>87) and GABA-d2 (106>89), were monitored in positive ionization mode. GABA was identified by comparing both mass spectra and retention time with those of pure standards injected in the same conditions. Peak area of GABA was normalized to internal standard area (GABA-d2) and sample dry weight. All experiments were repeated at least three times.

### Statistical analysis

Results are presented as the mean ± SD. Statistical analysis were performed by two-way ANOVA followed by a Tukey post hoc test when a significant difference was detected (different letters denote statistical significance at p < 0.05), or by two-tailed Student’s t-test (asterisks denote statistical significance at p < 0.05). Principal component analysis (PCA) was performed by means of the SIMCA version 13.0.3.0 software, using the log_2_ transformed data and unit variance normalization.

## ACKNOWLEDGMENTS

Metabolite measurements were carried out at Instituto de Biología Molecular y Celular de Plantas, CSIC-Universidad Politécnica de Valencia, and Servei Central d’Instrumentació Científica of the Universitat Jaume I.

## SUPPLEMENTAL MATERIAL

**Table S1.** Levels of metabolites involved in glycolysis, TCA cycle, and amino acid metabolism in Col-0 plants subjected to high light (HL), heat stress (HS) and the combination of HL and HS (HL+HS). Metabolite levels are expressed as the fold change compared to control conditions. *Abbreviations used*: CT, control; HL, high light; HS, heat stress; HL+HS, a combination of high light and heat stress.

**Table S2**. Expression level of transcripts involved in TCA cycle in Col-0 plants subjected to high light (HL), heat stress (HS) and the combination of HL and HS (HL+HS). Significant transcript levels (p < 0.05) are expressed as the fold change compared to control conditions. Data was obtained from the RNA-Seq analysis conducted by Balfagón et al. (2019). *Abbreviations used*: CT, control; HL, high light; HS, heat stress; HL+HS, a combination of high light and heat stress; n.s., not significant.

**Table S3.** Level of metabolites involved in glutamate metabolism in Col-0 plants subjected to high light (HL), heat stress (HS) and the combination of HL and HS (HL+HS). Metabolite levels are expressed as the fold change compared to control conditions. *Abbreviations used*: CT, control; GABA, γ-aminobutyric acid; HL, high light; HS, heat stress; HL+HS, a combination of high light and heat stress.

**Table S4.** Expression level of transcripts involved in GABA metabolism in Col-0 plants subjected to high light (HL), heat stress (HS) and the combination of HL and HS (HL+HS). Significant transcript levels (p < 0.05) are expressed as the fold change compared to control conditions. Data was obtained from the RNA-Seq analysis conducted by Balfagón et al. (2019). *Abbreviations used*: ALDH5F1, aldehyde dehydrogenase 5F1; CT, control; GAD, glutamate decarboxylase; HL, high light; HS, heat stress; HL+HS, a combination of high light and heat stress; n.s., not significant; POP2, pollen-pistil incompatibility 2.

**Fig. S1.** The experimental design used for the metabolomic study of high light (HL, yellow), heat stress (HS, orange) and a combination of high light and heat stress (HL+HS, grey) using Arabidopsis plants. HL was applied by exposing 30-day-old plants to 600 μmol m^−2^ s^−1^ (Philips, F54T5/TL84/HO/ALTO) at 23°C. HS was applied by transferring 30-day-old plants to 42°C. HL+HS was performed by simultaneously subjecting plants to 600 μmol m^−2^ s^−1^ and 42°C. Stress treatments were performed in parallel during 7 h. Following the stress treatments, plants were sampled for metabolomic analysis and gas exchange parameters were recorded. Another group of plants was allowed to recover under controlled conditions until flowering time to score for survival. 24 hours following the stress treatments, Leaf Damage Index (LDI) was also determined. All experiments were carried out at the same time-of-day during the light cycle (from 9 AM to 4 PM) and were repeated at least three times using Col-0 and *gad3* plants. *Abbreviations used*: CT, control; HL, high light; HS, heat stress; HL+HS, a combination of high light and heat stress; LDI, Leaf Damage Index.

**Fig. S2.** Enrichment of autophagy-related transcripts in the response of Arabidopsis plants to a combination of high light and heat stress. (A) Venn diagrams depicting the overlap between transcripts altered in Col-0 plants in response to a combination of high light and heat stress (HL+HS) and transcripts related to autophagy. (B) Heat map showing the expression levels of transcripts involved in autophagy in Col-0 plants subjected HL, HS and HL+HS combination. Non-significant expression levels compared to controls are shown in grey. Data was obtained from the RNA-Seq analysis conducted by Balfagón et al. (2019). *Abbreviations used*: HL, high light; HS, heat stress; HL+HS, a combination of high light and heat stress.

