## Supplemental Figures for "GABA plays a key role in plant acclimation to a combination of high light and heat stress"

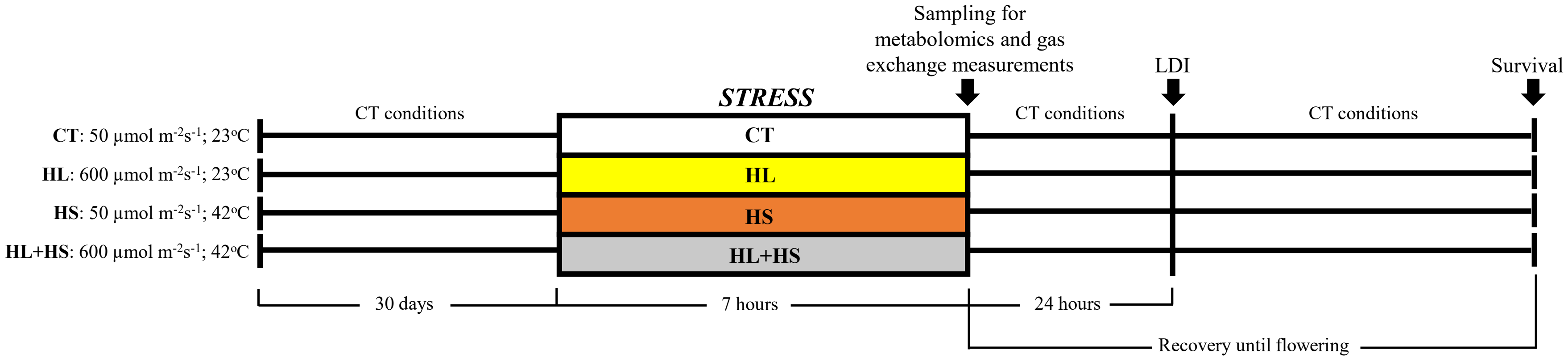


**Fig S1.** The experimental design used for the metabolomic study of high light (HL, yellow), heat stress (HS, orange) and a combination of high light and heat stress (HL+HS, grey) using Arabidopsis plants. HL was applied by exposing 30-day-old plants to 600 µmol $m^{-2}$ $s^{-1}$ (Philips, F54T5/TL84/HO/ALTO) at 23°C. HS was applied by transferring 30-day-old plants to 42°C. HL+HS was performed by simultaneously subjecting plants to 600 µmol $m^{-2}$ $s^{-1}$ and 42°C. Stress treatments were performed in parallel during 7 h. Following the stress treatments, plants were sampled for metabolomic analysis and gas exchange parameters were recorded. Another group of plants was allowed to recover under controlled conditions until flowering time to score for survival. 24 hours following the stress treatments, Leaf Damage Index (LDI) was also determined. All experiments were carried out at the same time-of-day during the light cycle (from 9 AM to 4 PM) and were repeated at least three times using Col-0 and *gad3* plants. *Abbreviations used*: CT, control; HL, high light; HS, heat stress; HL+HS, a combination of high light and heat stress; LDI, Leaf Damage Index.

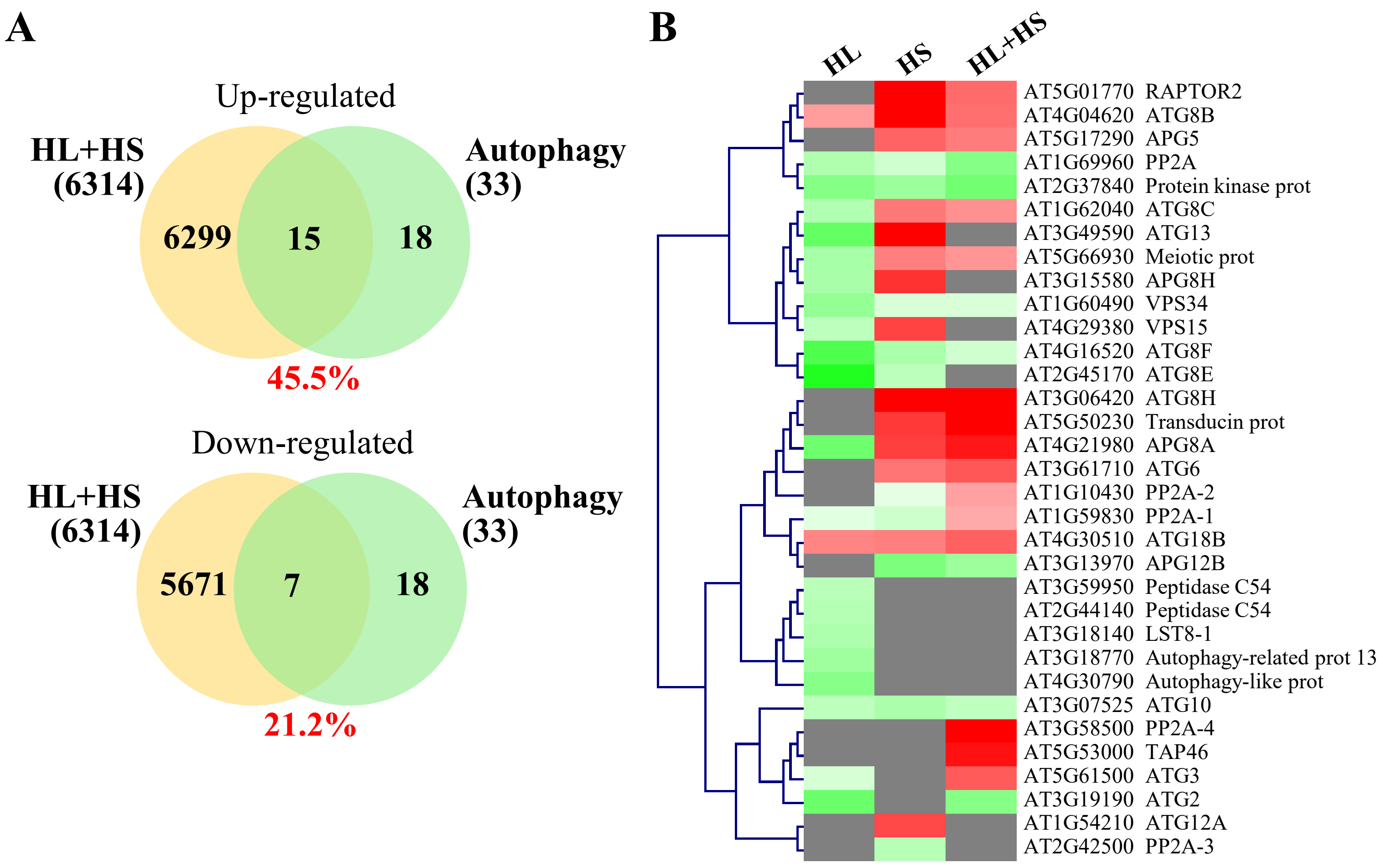


**Fig S2.** Enrichment of autophagy-related transcripts in the response of Arabidopsis plants to a combination of high light and heat stress. (A) Venn diagrams depicting the overlap between transcripts altered in Col-0 plants in response to a combination of high light and heat stress (HL+HS) and transcripts related to autophagy. (B) Heat map showing the expression levels of transcripts involved in autophagy in Col-0 plants subjected HL, HS and HL+HS combination. Non-significant expression levels compared to controls are shown in grey. Data was obtained from the RNA-Seq analysis conducted by Balfagón et al. (2019). *Abbreviations used*: HL, high light; HS, heat stress; HL+HS, a combination of high light and heat stress.
